# Slowing the Body slows down Time (Perception)

**DOI:** 10.1101/2020.10.26.355396

**Authors:** Rose De Kock, Weiwei Zhou, Wilsaan Mychal Joiner, Martin Wiener

## Abstract

Interval timing is a fundamental component action, and is susceptible to motor-related temporal distortions. Previous studies have shown that movement biases temporal estimates, but have primarily considered self-modulated movement only. However, real-world encounters often include situations in which movement is restricted or perturbed by environmental factors. In the following experiments, we introduced viscous movement environments to externally modulate movement and investigated the resulting effects on temporal perception. In two separate tasks, participants timed auditory intervals while moving a robotic arm that randomly applied four levels of viscosity. Results demonstrated that higher viscosity led to shorter perceived durations. Using a drift-diffusion model and a Bayesian observer model, we confirmed these biasing effects arose from perceptual mechanisms, instead of biases in decision making. These findings suggest that environmental perturbations are an important factor in movement-related temporal distortions, and enhance the current understanding of the interactions of motor activity and cognitive processes.

## Introduction

Interval timing is an essential part of survival for organisms living in an environment with rich temporal dynamics. Biologically-relevant behaviors often require precise calibration and execution of timed output on multiple levels of organization in the nervous system (Cisek and Kalaska, 2010). For example, central pattern generators produce basic locomotion in many organisms and yield balanced, rhythmic motor output via the oscillatory properties of inhibitory interneurons (Guertin, 2009). At greater levels of complexity, many behaviors rely on the explicit awareness of time (Buhusi and Meck, 2005). Subjective time is not always veridical, however; in fact, across many organisms, it is subject to distortion (Malapani and Fairhurst, 2002). As described by (Matthews and Meck, 2016), temporal distortions can arise from changes in perception, attention, and memory processes, and are proposed to be directly related to the vividness and ease of representation of a timed event. Interestingly, action properties can also influence perceived time. For example, it has been shown that subjective time on the scale of milliseconds to seconds is influenced by movement length (Yon et al., 2017), speed (Yokosaka et al., 2015), and direction (Tomassini and Morrone, 2016). More specifically, timed events accompanied by arm movements that are short (Yon et al., 2017), rapid (Yokosaka et al., 2015), or directed towards the body (Tomassini and Morrone, 2016) undergo compression.

These studies grant insight into the importance of action in the context of timing, but they are limited by focusing solely on volitional modulation of movement parameters. Often, organisms encounter changes in the environment that dramatically affect the way motor plans are executed. When these perturbations are encountered, organisms use feedback information to update current and future movement plans (Shadmehr et al., 2010). In the following experiments, we sought to modulate the parameters of movement length and speed by introducing changes in the movement environment itself rather than through instruction or task demands. Participants were required to time auditory tone intervals while moving a robotic arm manipulandum through environments with varying degrees of viscosity. This was tested first in a temporal bisection task, then in a temporal reproduction task with a new group of participants. If it is the case that time perception is biased by movement length (distance traveled during a timed event), then limiting movement by applying viscosity should lead to underestimation of intervals. In our previous work (Wiener et al., 2019), we utilized a very similar free-movement bisection paradigm to study the effect of movement on time perception. Unlike the current study, this paradigm had no viscosity factor, but rather tested whether participants that were allowed to move during timed intervals differed in performance from participants that were not allowed to move. Allowance of movement enhanced temporal perception by reducing variability (i.e., lower coefficient of variation). However, results observed in this study were mechanistically ambiguous. That is, we observed that temporal judgments were more precise with movement, but it was unclear whether this effect was driven by perceptual changes or modulation of decision properties (Figure 1a). Thus, in the current study we sought to provide a more mechanistic explanation of our observed results by disentangling perceptual effects and ensuing downstream processes (choice selection in the bisection experiment, and measurement and estimation in the reproduction experiment). In the bisection experiment, we demonstrate that viscosity successfully decreased movement length, and that this decrease was associated with underestimation of time intervals. We verified that this modulation was a result of interval timing and not a decision-related bias by applying a drift-diffusion model. In the reproduction experiment, all participants tended to overestimate durations, but viscosity was related to decreased overestimation and greater central tendency. We utilized a Bayesian Observer Model (Jazayeri and Shadlen, 2010; Remington et al., 2018) to verify that this effect was a result of perceptual bias rather than increased noise in the measurement and production processes. Overall, these results suggest that movement length has a direct influence on perceived interval length, regardless of whether this parameter is modulated by volitional or environmental factors.

**Figure 1:**
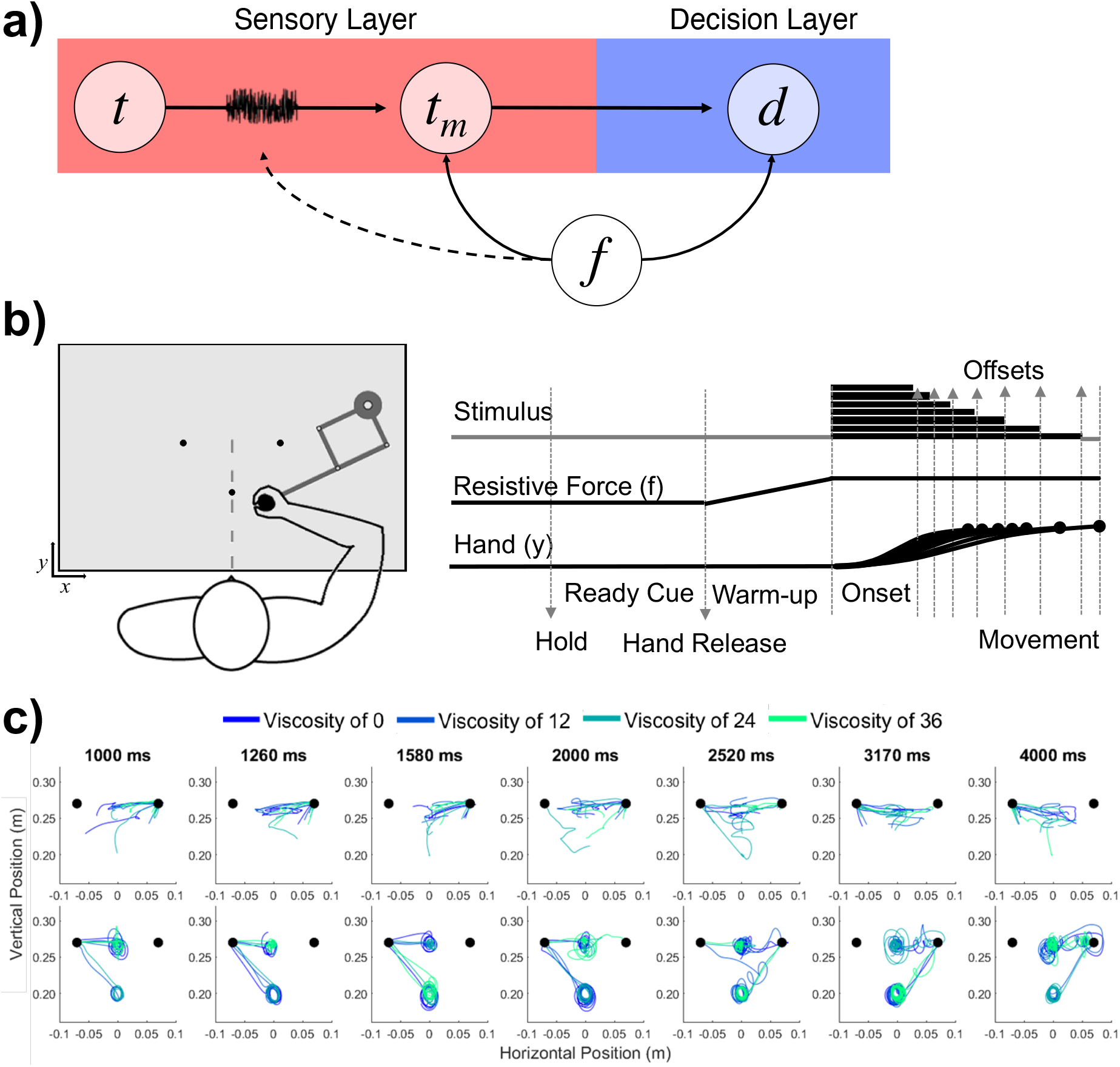
Hypothesis and Design. **a)** Potential pathways in which movement (*f*) could influences timing. The first possibility is that *f* specifically alters the perceptual layer, in which a stimulus presented for an amount of time (*t*) is perceived with noise as a temporal estimate (*t*_*m*_); here, *f* could specifically alter the measurement process, either by shifting the way that estimate is perceived or by altering the level of noise directly (dashed line). The second possibility is that *f* shifts the decision layer, such that decisions about time (*d*) are biased to one choice or another (e.g. more likely to choose “long”). **b)** Task schematic of Experiment 1. Participants began each trial with the robotic handle locked in a centralized location. The trial was initiated by a warm-up phase in which the hold was released and viscosity was applied in a ramping fashion until the target viscosity was reached. Participants were allowed to move throughout the workspace during warm-up and tone presentation, and reach to one of two choice targets to indicate their response. **c)** Each row displays sample trajectories from two subjects. The trajectories include movement during the tone and the ensuing decision for the seven possible tone durations.

## Results

### Experiment 1 - Temporal Bisection

In our first experiment, 28 human subjects engaged in an auditory temporal bisection task using supra-second intervals between 1-4s. Subjects were required to classify each interval as “long” or “short”, compared to the running average of all previously experienced intervals. To classify each interval, subjects were required to move the arm of a robotic manipulandum to one of two response locations, counterbalanced between subjects. Prior to tone onset, subjects were allowed a 2s “warm-up” period, in which they were free to move the cursor around and explore the environment. During this period, the resistive force (f) against the manipulandum was gradually increased to reach a peak viscosity (v) of four possible levels (0, 12, 24, or 36 Ns/m^2^; see Materials and Methods); the viscosity remained at this level for the duration of the tone, returning to zero at the interval offset (Figure 1b). Entry into the response location prior to the tone offset was penalized by restarting the trial, and so the optimal strategy was to move the cursor closer to the “short” location, and then gradually move to the “long” location as the tone elapses (Wiener et al., 2019). Consistent with this strategy, we found that the relative location of the hand at interval offset was closer to the short target for intervals at or under the middle of the stimulus set, but rapidly moved closer to the long target for longer intervals [*F* (6,162)=4.791, *p* < 0.001, *η*^2^_p_=0.151] (Figure 2e). However, no impact of viscosity was observed on relative hand position [F(3,81)=1.595, p=0.197], suggesting that movement had little impact on the ability of subjects to employ this strategy (Figure 1c). We further verified that our viscosity manipulation worked, by observing a decrease in movement lengths [F(3,81)=21.05, p < 0.001, \etaη2p=0.438] and an increase in force applied to the arm with higher viscosities [F(3,81)=22.736, p < 0.001, *η*^2^_p_ =0.457] (Figure 2f &g).

**Figure 2:**
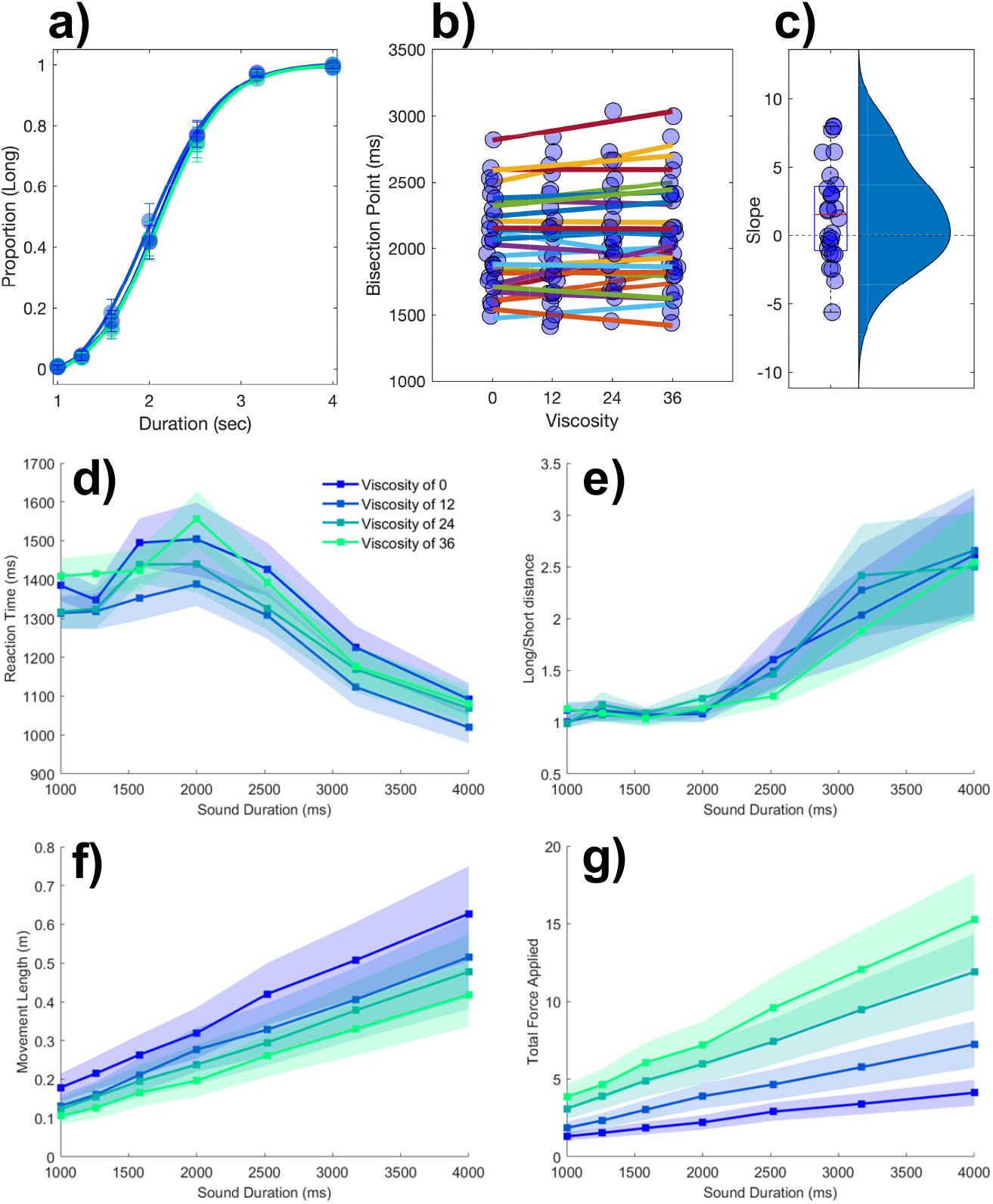
Viscosity shifts time responses. **a)** Group psychometric functions separated by viscosity. **b)** Individual bisection point data by viscosity and **c)** associated slopes when fitted with a linear function. **d)** Mean reaction times, **e)** Euclidean distance ratio to long and short targets at tone offset, **f)** movement length (during tone), and **g)** cumulative force applied (during tone). Shaded error bars represent the standard error of the mean.

Analysis of choice responses proceeded by constructing psychometric curves from the mean proportion of “long” response choices for each interval/viscosity combination, and chronometric curves from the mean reaction time (RT) as well (Figure 2a & d). Psychometric curves were additionally fit with cumulative Gumbel distributions, from which the bisection point (BP) was determined as the 0.5 probability of classifying an interval as long. Analysis of the BP values across all four viscosities with a repeated-measures ANOVA revealed a significant effect of viscosity [*F* (3,81)=3.774, *p*=0.014, *η*^2^_p_ = 0.123] (Figure 2b). A further examination revealed this to be a linear effect, with BP values generally increasing with viscosity, indicating a greater tendency to classify intervals as “short” [*F* (1,27)=5.439, *p* = 0.027, *η*^2^_p_ = 0.168] (Figure 2c); we further note that examination of a quadratic contrast did not reveal a significant effect [*F* (1,27)=1.209, *p*=0.281]. Analysis of RT values demonstrated faster RTs with longer perceived duration, consistent with previous reports. Additionally, a significant effect of viscosity was observed [F(3,81)=38.302, p < 0.001, *η*^2^_p_ = 0.587]; however, the effect was variable across viscosities, with faster RTs for mid-range viscosities. No effect of viscosity was observed on the CV [F(3,81)=0.377, p=0.77].

In our previous report (Wiener et al., 2019), we observed that subjects performing this task exhibited a more precise perception (lower CV) of time compared to subjects who performed a different version where the robot arm was fixed at the starting point for the duration of the interval. Although the present study allowed all subjects to move freely during the interval, we hypothesized that movement during the interval would interact with precision within-subject. To test this possibility, we performed a median split of the movement length for each interval/viscosity combination and re-analyzed the psychometric curves for each viscosity condition. For the BP, we again observed an increase with viscosity [*F* (1,27)=5.936, *p* = 0.022, *η*^2^_p_=0.18], regardless of how much subjects moved [*F* (1,27)=1.397, *p*=0.247] (Figure 3a). For the CV, a significant interaction between viscosity and movement length was observed [*F* (1,27)=7.694, *p*=0.01, *η*^2^_p_=0.222], in which the CV was significantly lower when subjects moved more, but only when the viscosity was zero [*t* (27)=−2.237, *p*=0.034, *D*=1.2] (Figure 3b), and not for any other viscosity (all *p*>0.05). This indicates that precision again improved with greater movement, but only when no impediments from a viscous movement environment existed.

**Figure 3:**
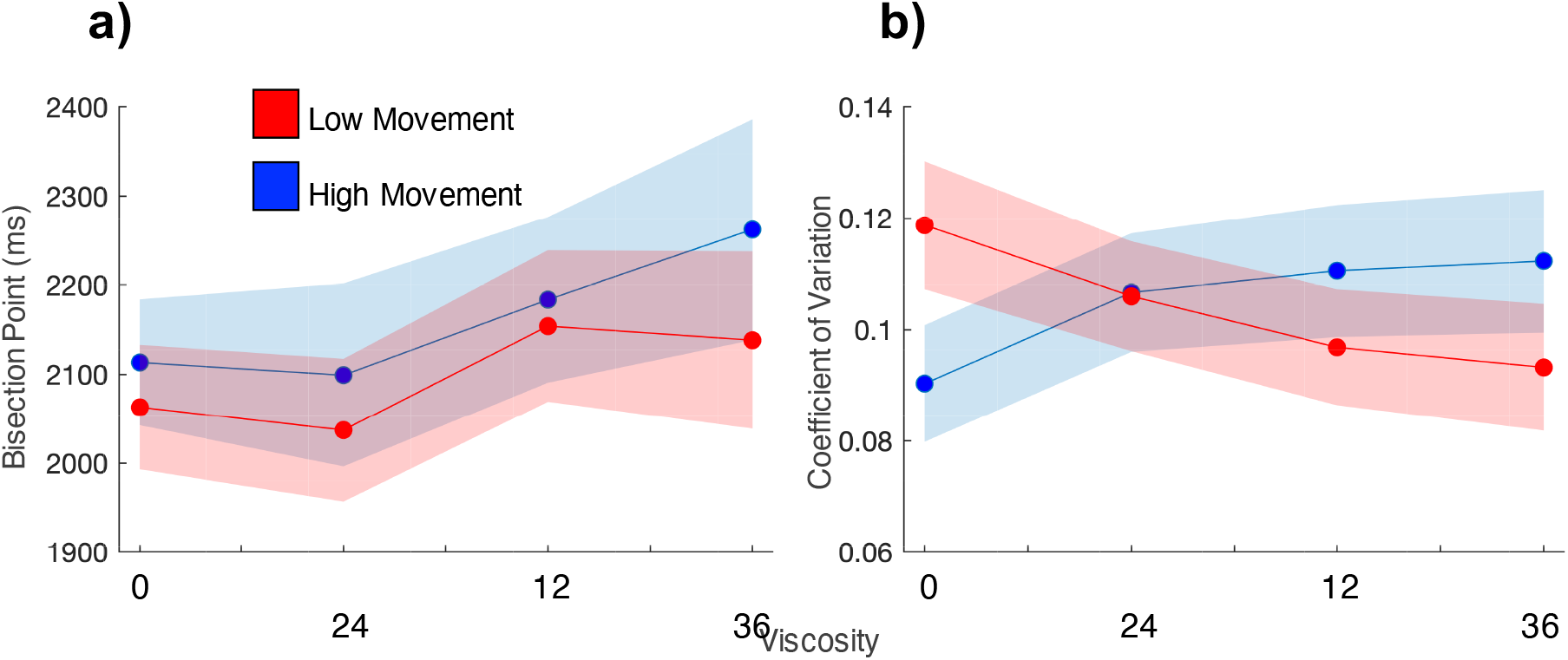
Movement speed does not change viscosity effects, but does influence precision. **a)** Mean bisection points and **b)** coefficients of variation for participant trials divided into high and low movement via a median split. Viscosity shifted the bisection point across both movement types (left); however, precision was influenced by movement, but only when no viscosity existed.

#### Drift Diffusion Modeling

The results of this Experiment appeared to support the hypothesis that increasing viscosity while judging an auditory interval led to a shorter perception of that interval. However, as stated in the Introduction, a shift resulting from increased viscosity could have either altered perception or biased subjects to classify intervals as “short”; both outcomes could explain our results, as our task inherently involves a directional judgment (Yates et al., 2012; Schneider and Komlos, 2008). To further tease apart these two possibilities, we decomposed choice and RT data using a drift diffusion model (DDM) of perception and decision making. We employed hierarchical DDM (HDDM; (Wiecki et al., 2013)) in order to constrain fitted parameters for individual subjects by the group mean (see Materials and Methods). Under this framework, evidence is accumulated over time towards a “long” or “short” decision boundary. A shift in the drift rate towards one of these boundaries is interpreted as evidence in favor of a shift in the perceptual evidence, whereas a change in either the threshold boundary or the starting point could be interpreted as a change in the decision layer (Voss et al., 2004).

In our analysis of the fitted DDM parameters, we observed first a significant shift in the drift rate (*v*) with increases in viscosity [F(3,81)=4.562, p=0.005, *η*^2^_p_ =0.145]; this shift was notably linear in nature, with the drift rate shifting linearly to the “short” duration boundary with higher viscosities [F(1,27)=7.866, p = 0.009, *η*^2^_p_ =0.226]. Further analyses also revealed significant effects of viscosity on the threshold (*a*) [F(3,81)=12.356, p <0.001, *η*^2^_p_ =0.314] and starting point (z) [F(3,81)=43.73, p <0.001, *η*^2^_p_ =0.618], but no effect on the non-decision time [F(3,81)=2.257, p =0.088] (Figure 4b). In both cases for the threshold and starting point, the dominant pattern was for these values to drop for viscosities above zero, but show little variation beyond that. Given the linear pattern observed for changes in the drift rate, we further explored whether this parameter could exclusively explain the shift in the BP. To test this, we calculated the slope of a linear regression for each parameter against viscosity for each subject, and correlated these with the slope values for BP against viscosity. Here, the only significant correlation observed was for the drift rate [Pearson = −0.5132, *p* = 0.0052; Spearman = −0.7865, *p*<0.001], and not for any other parameter (all *p*>0.05). A Fisher’s Z-test comparing this correlation confirmed that it was significantly greater than for all other parameters (see Table 1).

**Figure 4:**
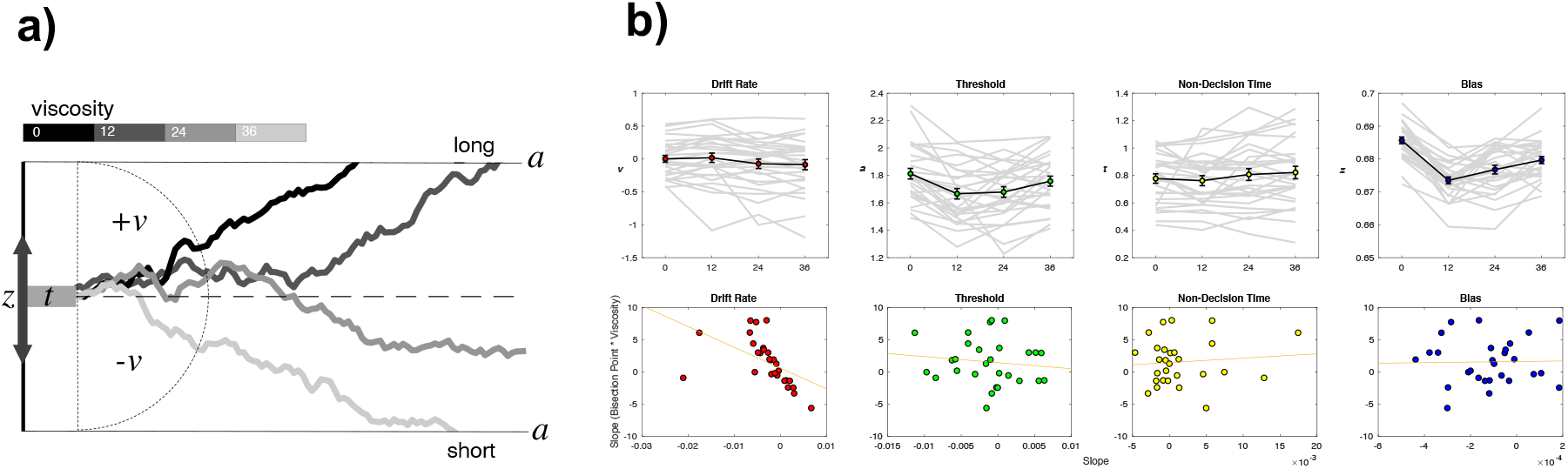
Drift Diffusion Modeling of Bisection Performance and Viscosity. **a)** Example DDM model, in which evidence is accumulated to one of two decision bounds (“long” and “short”), separated by *a*. Evidence accumulation drifts at particular rate (*v*) that can be positive or negative, depending on the direction of the drift to a particular boundary. The drift rate is additionally delayed by non-decision time (*t*) and may be biased towards one of the boundaries by a certain amount (*z*). Viscosity was specifically found to influence the drift rate, which higher viscosities associated with a shift in drift from the long to short decision boundary (presented traces represent example simulations). **b)** Fitted DDM results for all four parameters, showing that viscosity linearly shifted the drift rate, but also modulated threshold and bias parameters in a nonlinear (stepwise) manner. Below panels demonstrate the correlation between DDM parameters and the viscosity effect on behavior; only drift rate exhibited a significant correlation (see also Table 1 for Fisher Z comparisons between correlations). Left panel was additionally significant after outlier removal.

**Figure 5:**
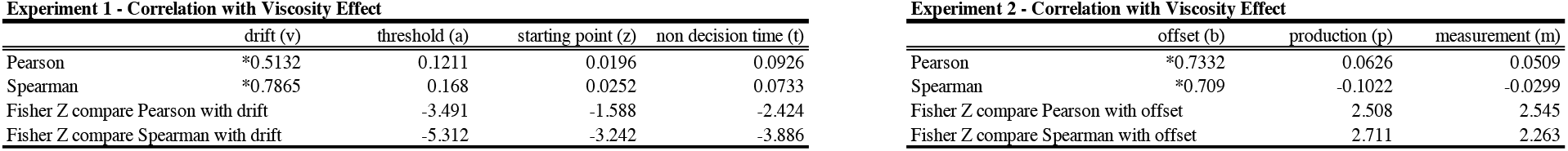
**Table 1** - Correlation coefficients and Fisher Z comparisons between fitted parameters and behavioral effects.

### Experiment 2 - Temporal Reproduction

The results of Experiment 1 demonstrated that increased resistive force while subjects made temporal judgments about auditory durations led to shorter reported lengths of those durations Computational modeling using a DDM further suggested that this shift was due to viscosity altering the perceived duration, rather than altering decision bias. However, in this experiment, decision-making and perception are intertwined, such that subjects must simultaneously measure the interval duration while classifying it. Indeed, previous research has suggested that, once the categorical boundary (here, the BP) has been crossed, subjects stop accumulating temporal information (Wiener and Thompson, 2015).

To further disentangle whether viscosity impacts perception or decision layers, we had a new set of subjects (n=18) perform a temporal reproduction task, in which they moved the robotic arm while listening to auditory tone intervals and encoding their duration (Figure 6). Following this, the arm was locked in place and subjects reproduced the duration via a button-press attached to the handle (see Materials and Methods). Viscosity was again randomized across the same four levels during the encoding phase; in this way, the impact of resistive force was applied only while subjects were actively perceiving duration, without any deliberative process.

**Figure 6:**
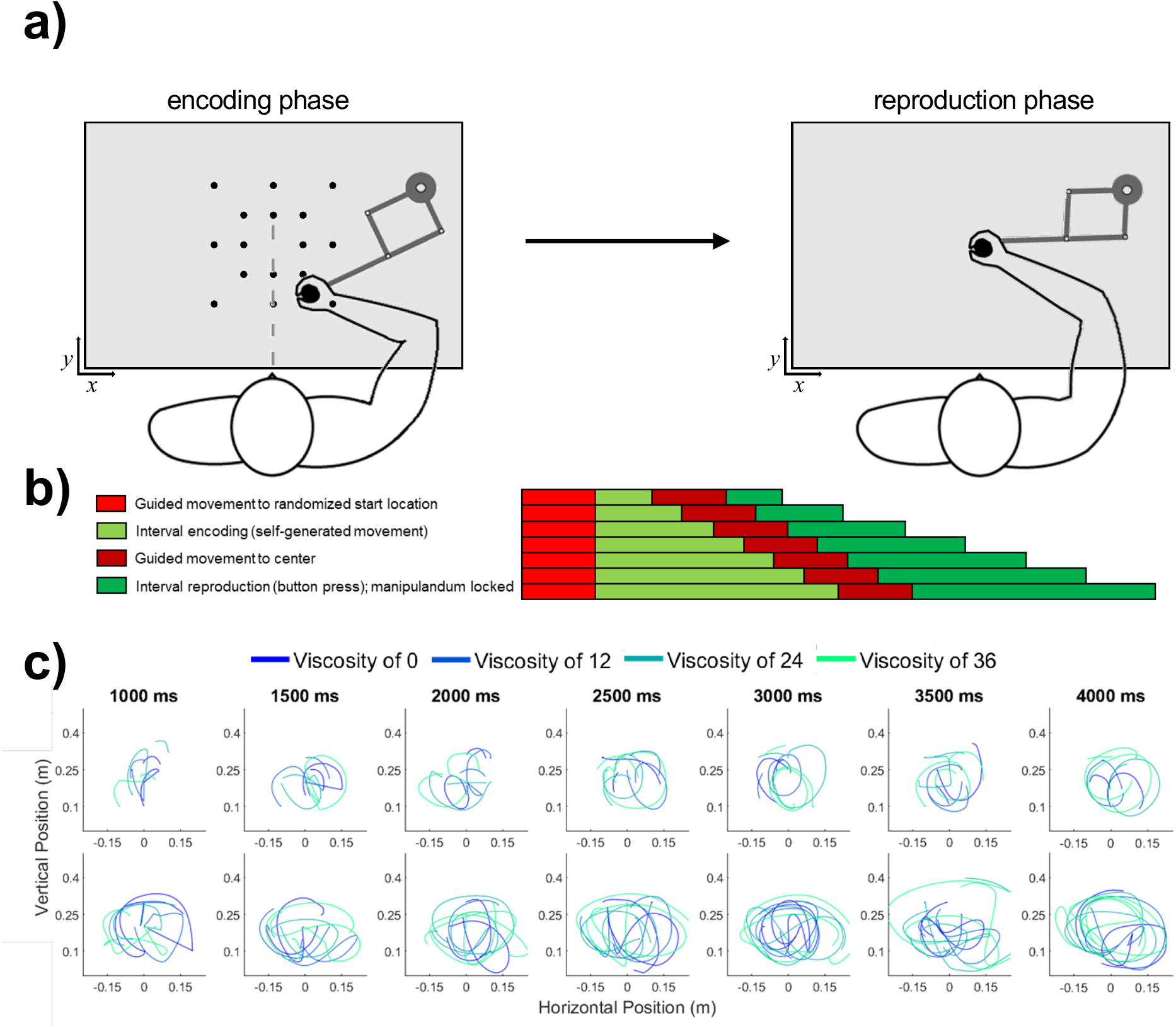
Task schematic of Experiment 2. **a)** Participants began each trial at a randomized start location and were required to initiate movement in order for the test duration to play (encoding phase). Unlike Experiment 1, the desired viscosity was applied immediately rather than in a ramping fashion. Then, the handle was brought to a central location where participants reproduced the duration by holding and releasing a button attached to the handle. **b)** Timeline for each of the seven tested intervals. **c)** As in Experiment 1, each row displays sample trajectories from two subjects for the seven possible tone durations.

We initially confirmed again that our viscosity manipulation was effective, with reduced movement length [F(3,51)=149.82, p < 0.001, *η*^2^_p_ =0.898] and increased force [F(3,51)=114.84, p < 0.001, *η*^2^_p_ =0.871] observed with greater viscosities (Figure 7e & f). For behavioral results, we initially measured the reproduced durations (t_p_), finding both a main effect of viscosity [F(3,51)=5.5, p = 0.002, *η*^2^_p_ =0.244] and an interaction with the presented duration [F(3,51)=1.814, p = 0.023, *η*^2^_p_ =0.096] (Figure 7c). No impact on the variance of reproduced estimates was observed, with the CV remaining stable across all viscosities [F(3,51)=0.691, p = 0.562] (Figure 7d). More specifically, we observed that reproduced durations generally were overestimated compared to the presented sample durations (t_s_), this effect was quantified by measuring the offset for each reproduced duration compared to the presented one; here, we additionally observed an effect of viscosity, with less overestimation with increasing viscosity [F(3,51)=5.5, p = 0.002, *η*^2^_p_ =0.244] (Figure 7a & b). We additionally observed an increase in the so-called central tendency effect, in which reproduced durations gravitate to the mean of the stimulus set, with greater viscosities; this effect was quantified by a change in slope values of a simple linear regression [F(3,51)=3.473, p = 0.023, *η*^2^_p_ =0.17].

**Figure 7:**
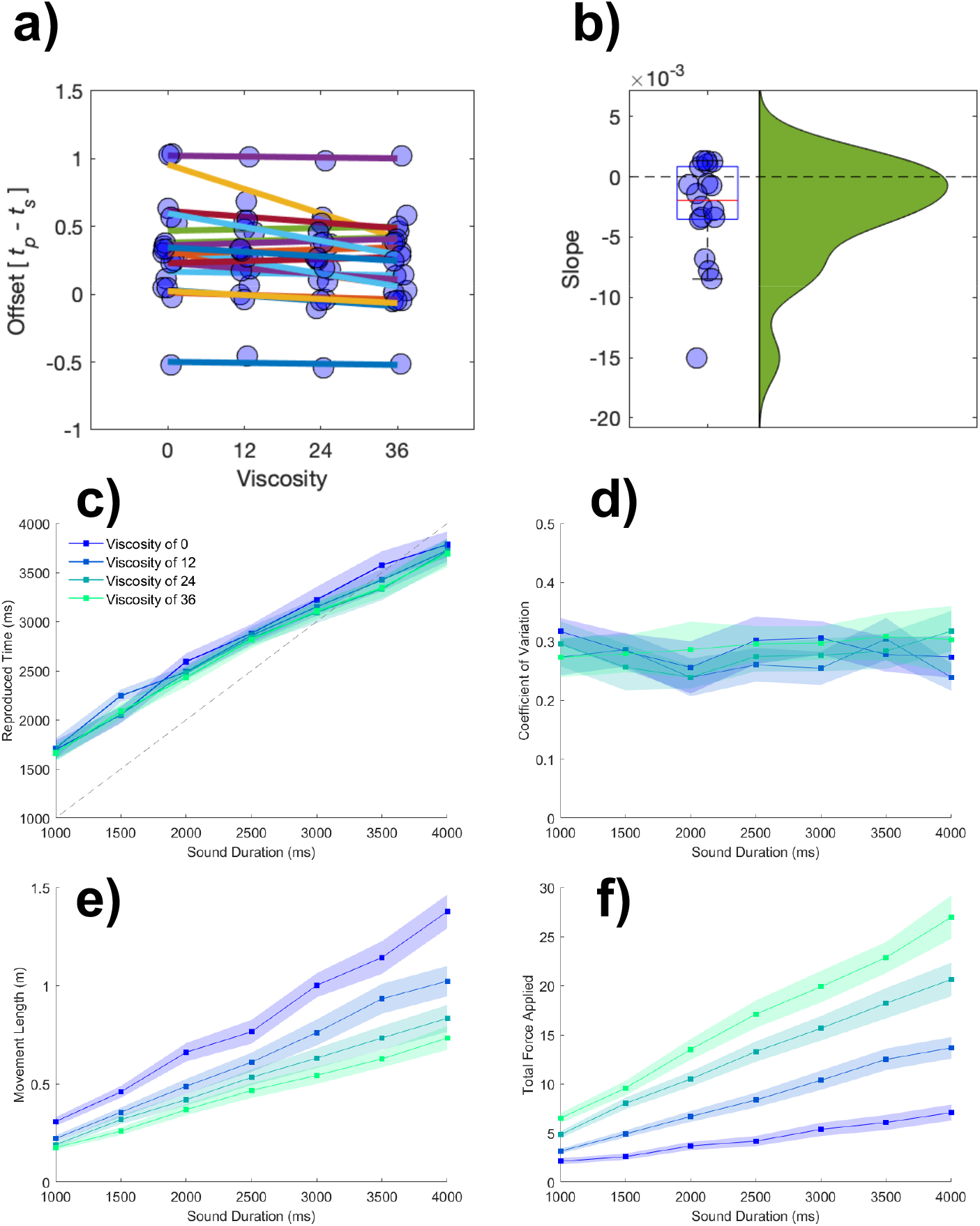
Viscosity shifts time reproduction. **a)** Mean offset (t_p_ - t_s_) across duration for each viscosity condition. Greater viscosity led to a decrease in offset (shorter reproduced durations). **b)** Individual slope values with distribution. **c)** Mean reproduction times, **d)** coefficient of variation, **e)** movement length (during tone), and **f)** cumulative force applied (during tone). Shaded error bars represent the standard error of the mean.

#### Bayesian Observer Model

The results of Experiment 2 revealed that, with increasing viscosity while encoding a time interval, the reproduced interval was increasingly, relatively, shorter in length. Again, this finding is consistent with reduced movement altering the perception of temporal intervals. We note that the temporal reproduction task as designed does not share the overlap with decision-making as in the temporal bisection task, as viscosity was only manipulated while subjects estimated the interval, and was not included during reproduction. However, we also note that the behavioral data alone are somewhat ambiguous to how viscosity impacts time estimation, as we observed both a shift in time intervals, as well as an increase in central tendency with greater viscosities. Changes in central tendency may be ascribed to a shift in uncertainty while estimating intervals, and although the CV did not change across viscosities, it remains possible that viscosity led to greater uncertainty, which would explain the observed shifts.

To tease these two possibilities apart, we employed a Bayesian Observer-Actor Model previously described by Remington and colleagues (Remington et al., 2018; Jazayeri and Shadlen, 2010) (see methods). In this model, sample durations (t_s_) are inferred as draws from noisy measurement distributions (t_m_) that scale in width according to the length of the presented interval. These measurements, when perceived, may be offset from veridical estimates as a result of perceptual bias or other outside forces (b). Due to the noise in the measurement process, the brain combines the perceived measurement with the prior distribution of presented intervals in a statistically optimal manner to produce a posterior estimate of time. The mean of the posterior distribution is then, in turn, used to guide the reproduced interval (t_p_), corrupted by production noise (p) (Figure 8a). The resulting fits to this model thus produce an estimate of the measurement noise (m), the production noise (p), and the offset shift in perceived duration (b). Note that the offset term is also similar to that employed for other reproduction tasks as a shift parameter (Petzschner and Glasauer, 2011).

**Figure 8:**
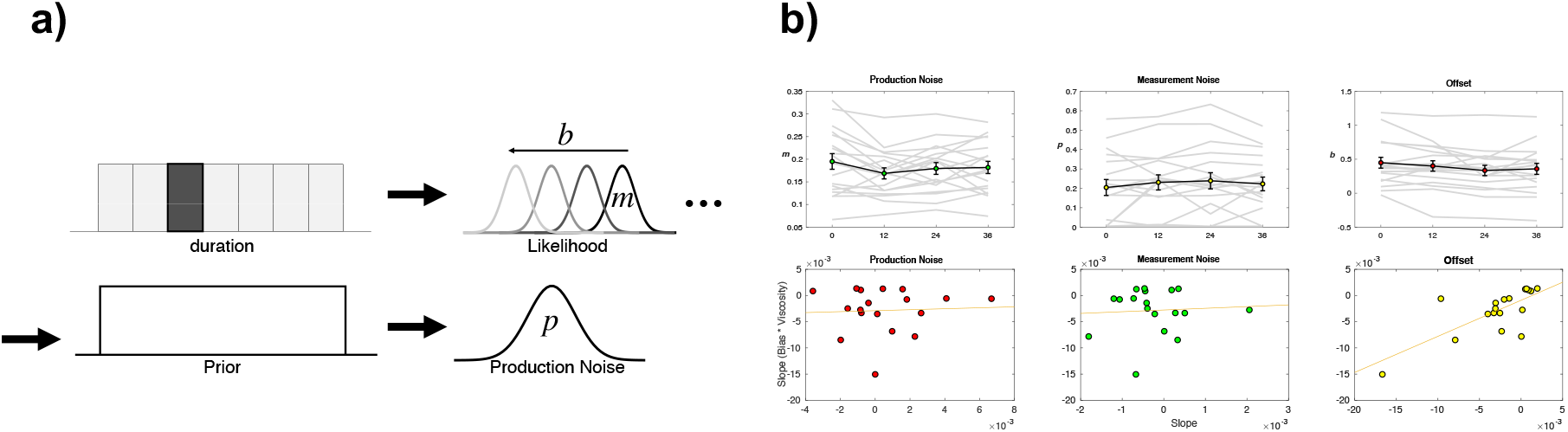
Bayesian Observer Model of Reproduction Performance and Viscosity. **a)** Example Observer Model. On a given trial, a presented duration is drawn from a likelihood distribution with scalar variance leading to a measurement estimate (*m*) that is shifted by an offset parameter (*b*). The measurement estimate is combined with a uniform prior distribution of presented durations, and then finally affected by production noise (*p*). Viscosity was found to specifically shift *b* in a linear manner, with greater viscosities associated with shorter perceived durations. **b)** Fitted results for all three parameters, demonstrating a linear effect of offset, no effect of measurement noise, and a nonlinear (stepwise) shift in production noise with greater viscosities. Bottom panels display correlations with the behavioral effect of viscosity; only the offset parameters exhibited a significant effect (see Table 1 for Fisher Z comparisons). Right panel was additionally significant after outlier removal.

The result of the model fitting first demonstrated a significant effect on the width of the production noise (p) [F(3,51)=3.548, p = 0.021, *η*^2^_p_ =0.173] (Figure 8b). More specifically, production noise was found to decrease with higher viscosities; however, this effect was not linear, with the only difference being for zero viscosity estimates higher than all others. For the offset shift (b), we observed a significant effect of viscosity [F(3,51)=3.72, p = 0.017, *η*^2^_p_ =0.18] that was linear in nature, with a reduction in values with increasing viscosity. No effect of viscosity was observed on the measurement noise parameter (m) [F(3,51)=1.212, p = 0.315]. As with the DDM results of Experiment 1, we further explored whether the linear nature of the shift in *b* could best explain the observed underestimation of duration by calculating the slope of a regression line for each parameter against viscosity and compared that to the change in reproduced duration. Only the offset term significantly correlated with the underestimation effect [Pearson = 0.7332, p < 001; Spearman = 0.709, p<0.001]. Again, a Fisher’s Z-test comparing this correlation confirmed that it was significantly greater than for all other parameters (see Table 1).

## Discussion

The above experiments demonstrate that systematically impeding movement during interval timing leads to a subsequent compression of perceived duration. These findings complement previous work showing that time perception is highly sensitive to movement ((Yon et al., 2017); (Yokosaka et al., 2015); (Tomassini and Morrone, 2016)), and here we confirm a case in which movement parameters (e.g., length and duration) did not have to be self-modulated to induce these distortions. The movement restrictions we implemented (i.e., moving in environments with different manipulations of viscosity) tended to shift the BP later in time in a temporal bisection task, and subsequently shortened perceived intervals in a temporal reproduction task.

In the temporal bisection task we found that increased viscosity, on average, shifted the BP such that subjects responded ‘long’ less often. We then applied a drift-diffusion model to isolate the cognitive mechanisms contributing to this effect (i.e., whether it was a function of decision bias, speed-accuracy trade-off calibration, non-decision time, or the rate of evidence accumulation; (Ratcliff, 1978)). The only significant contributor was the drift rate parameter, which linearly shifted from the ‘long’ to the ‘short’ boundary with increasing viscosity. While this was evidence for a purely perceptual effect of viscosity on perceived time, we further investigated this effect by administering a temporal reproduction task in Experiment 2. Eliminating the decision process ensured that recorded responses were more representative of timing distortions via perceptual modulation. Here, participants made temporal estimations during movement, and reproduced these via button press. Whereas in the temporal bisection task the degree of movement between participants was self-selected and highly variable, we attempted to reduce this variability by requiring participants to move continuously during trials in the temporal reproduction task. We also controlled for performance variability due to familiarization by including a brief training session. Participants exhibited an overestimation bias when reproducing the interval duration, a result previously shown in motor reproduction of auditory intervals (Shi et al., 2013). However, the degree of overestimation decreased as a function of viscosity, confirming the compression effect seen in the temporal bisection task. Using a Bayesian observer model ((Jazayeri and Shadlen, 2010); (Remington et al., 2018)), we observed a linear shift in perceptual bias as a function of viscosity, further supporting a purely perceptual effect of movement slowing on timing.

The results between experiments converge on the finding that viscosity manipulation leads to interval underestimation (reflected in the bias parameters). Further, timing precision was generally unaffected by our manipulations. In the temporal bisection task, the CV remained constant across viscosities; the only notable CV effect was revealed by the median split analysis, in which trials with greater movement led to greater precision for the zero viscosity condition. This was in agreement with our previous study demonstrating a movement-related enhancement in temporal precision estimates (Wiener et al., 2019), and suggests that free movement can improve timing precision only in unrestricted movement environments. The temporal reproduction task also showed that viscosity did not affect the variability of estimation (CV), but interestingly, may have been associated with greater uncertainty as indicated by increased central tendency of reproduction slopes.

In addition to the paradigm difference in Experiments 1 and 2, it is worthwhile to consider the methodological differences that may have influenced these results. As mentioned above, in our first experiment we allowed movement, while in the second experiment we required it. Additionally, movement in the temporal bisection task had utilitarian value; participants could strategically approach candidate targets, and thus movement offered the potential to improve performance by shortening RTs. Movement in the temporal reproduction task did not provide this decision-making advantage, but notably the perceptual biases due to viscosity maintained the same directionality across experiments. That is, in response to viscosity, the BP parameter in Experiment 1 and the perceptual bias parameter in Experiment 2 shifted upwards and downwards respectively, in accordance with temporal compression. This suggests that movement influenced similar temporal estimation mechanisms, and despite methodological differences, the biasing effect of viscous environments was robust under different task demands. One additional note regarding this work is that subjects were not required to time their movements but were rather using their movements for timing. We believe the distinction here is important, as the ancillary movement patterns nevertheless influenced the perceived timing.

Our results parallel prior work investigating temporal distortions as a function of movement parameters. (Press et al., 2014) presented tactile stimulation to participants’ fingers during movement or while stationary as they viewed congruent or incongruent hand avatars displayed on a screen. The duration of the tactile stimulus was perceived as longer when the avatar was congruent to concurrent movement. (Yon et al., 2017) similarly found that when participants executed finger movements of a pre-specified duration and listened to an auditory stimulus with an independently-selected duration, judgments were biased towards the duration of the movement. As in our experiments, these findings support that sensory estimates of duration are strongly biased by motor activity. In contrast, some prior accounts do not align with our results. For example, (Yokosaka et al., 2015) found that visual intervals demarcated by pairs of visual flashes were compressed during fast hand movement, whereas in our experiment we show that slowing down movement leads to compression. Additionally, Tomassini and colleagues (Tomassini et al., 2014) reported that tactile intervals were compressed during hand movement. Considering these examples, a crucial note–as highlighted in (Iwasaki et al., 2017)–is that many of the distortive effects of movement can be linked to whether an interval is filled or unfilled. Indeed, while we utilized filled auditory intervals in our tasks, the studies with contrasting effects utilized unfilled intervals. Also of interest is the type of movement in the listed studies and the interval ranges used. Movements were typically stereotyped across trials, and intervals were tested in the subsecond range. Here, we allowed participants to move freely along a two-dimensional plane and there were considerable individual differences in selected trajectories. Most notably, it was the *externally* imposed restriction of movement that turned out to be more influential on temporal perception than self-modulated movement characteristics. This novel contribution to the existing body of research highlights the importance of sensory feedback in timing, whereas the study of movement-induced time distortions has focused primarily on feedforward effects. These complementary accounts enrich the current understanding of the coupling of movement and time perception, and support the longstanding notion that interval timing in the brain utilizes multiple streams of sensory information and distributed timing circuits to form a unified percept of duration ((Chen and Vroomen, 2013); (Bausenhart et al., 2014); (Wiener et al., 2011)). Understanding the integration of these signals is an important problem in modern neuroscience, and here we have presented a strong case for greater investigation into the role of movement perturbations in time perception.

Beyond understanding mechanisms of temporal processing and movement, these approaches may be of interest in order to study clinical disorders for which timing and movement are disrupted. Notable work in recent years has strongly suggested that motor control is an extension of ongoing cognitive computations ((Lepora and Pezzulo, 2015); (Resulaj et al., 2009)), and adopting an integrated view of cognition-action pathways is a promising avenue for understanding these disorders and developing therapies that exploit these links. For example, in the case of Parkinson’s (PD) and Huntington’s Diseases (HD), core timing networks in the brain overlap substantially with the motor circuitry targeted by neural degeneration, such as the basal ganglia ((Obeso et al., 2000); (Browne et al., 1997)). Motor deficits in these diseases are often accompanied by timing deficits (Avanzino et al., 2016) and other cognitive abnormalities ((Robbins and Cools, 2014); (Paulsen, 2011)). The shared neural circuitry combined with these parallel deficits provide a basis for incorporating movement into cognitive training and vice versa.

In contrast, psychiatric and neurodevelopmental disorders are usually discussed in terms of cognitive deficits despite exhibiting motor idiosyncrasies. Although embodied cognition has gained traction in basic science research, there are fewer approaches in clinical research that consider cognitive and motor symptoms in relation to one another. One example is attention deficit hyperactivity disorder (ADHD). The excessive motoric activity (i.e., hyperactivity) associated with the disorder is typically seen as disruptive, but interestingly, some studies have shown that it can boost cognitive control performance ((Rapport et al., 2009); (Hartanto et al., 2016)). In light of the timing deficits present in ADHD (Plummer and Humphrey, 2009), it would be interesting to explore the extent movement can provide a similar benefit to timing performance. The coupling of timing and motor functions taken together with the supramodal nature of core timing circuits provides an excellent opportunity to probe cognition-action pathways in various clinical disorders. On the one hand, movement can sharpen certain perceptual and cognitive processes, and on the other hand, it can introduce biasing effects on timing and on other perceptual judgments (Moher and Song, 2014), including when movement perturbations are applied (Hagura et al., 2017). Therefore, further investigation is warranted to better understand how these effects can be exploited to improve outcomes for patients.

In summary, we tested participants’ ability to time intervals while moving, and in two separate timing paradigms demonstrated that imposing restrictions on movement subsequently shortened perceived time. Computational modeling confirmed that these effects arose from perceptual differences rather than downstream cognitive processes. The influence of motor activity on sensory processing is well studied–motor activity can modulate sensory processing across modalities, as early as in primary sensory cortices (for review, see (Parker et al., 2020)). For example, locomotion can modulate gain in rodent visual and auditory cortices ((Niell and Stryker, 2010); (Zhou et al., 2014)), and in humans, orientation change detection is improved when preparing to make a grasping motion aligned with the original orientation (Gutteling et al., 2011). Recent reports have shown that timing, through a neurally distributed system, is also modulated by and can even be improved by movement ((Wiener et al., 2019); (Carlini and French, 2014); (Manning and Schutz, 2013)). Our work confirms that time is subject to distortion by externally-imposed movement constraints and allowances, at least in the auditory modality and within the suprasecond range. It is important to note that timing distortions can be modality-specific (Bueti, 2011), and as stated above, differ when intervals are filled versus unfilled (Iwasaki et al., 2017). These are natural considerations regarding our results, and could form the basis for future work to investigate different modalities, temporal ranges, and interval presentation styles. Another consideration is that in our experiments, viscosity was fixed throughout each trial; in followup experiments, it would be interesting to calibrate the nature and degree of movement perturbations to observe the resulting effects on timing. For example, experimenters could introduce dynamic perturbations or alter visual feedback much like in motor adaptation experiments ((Shadmehr and Mussa-Ivaldi, 1994); (Krakauer et al., 2000); (Alhussein et al., 2019); (McKenna et al., 2017); (Zhou et al., 2017)).

## Materials and Methods

### Participants

28 participants took part in Experiment 1 (18 female, 10 male, M age = 23.5(7.0)) and 18 separate participants took part in Experiment 2 (7 female, 11 male, M age = 21.5(4.1)) for $15 per hour in gift card credit. These sample sizes were chosen to accord with our previous report (Wiener et al., 2019). All participants were right-handed as measured by the Edinburgh Handedness Inventory (Oldfield, 1971). All protocols were approved by the Institutional Review Board at the University of California, Davis.

### Apparatus

Both experiments utilized a robotic arm manipulandum (KINARM End-Point Lab, BKIN Technologies; (Nguyen et al., 2019); (Hosseini et al., 2017)). Here, the participants manipulated the right arm on a planar workspace to perform the tasks and were blocked from viewing their arm directly by a horizontal screen display. A downward-facing LCD monitor reflected by an upward-facing mirror allowed viewing of experiment start locations and targets, demarcated by small circles. Participants were seated in an adjustable chair so that they could comfortably view the mirrored display. In Experiment 1, a cursor on the screen projected their current arm position during each trial. The manipulandum sampled motor output at 1000 Hz.

### Procedure

#### Experiment 1

In the first experiment, participants performed an auditory temporal bisection procedure for intervals of 1000, 1260, 1580, 2000, 2520, 3170, and 4000 ms for a 440 Hz tone. A total of 280 trials were segmented into five blocks with the option for a short (1-2 min) break between each block. Participants were instructed to start each trial in a central target location, where the manipulandum locked the arm in place for 1000 ms. A warm-up phase began as the hold was released and the words “Get Ready” were displayed on the screen. Participants were encouraged during instruction to move freely throughout the workspace during each trial, and respond as quickly and accurately as possible. During the warm-up phase, viscosity was applied in a linearly ramping fashion, reaching one of four viscosity values (0, 12, 24, or 36 Ns/m^2^) in 2000 ms. Simultaneously, two response targets appeared at 105° and 75°, equidistant from the starting location. Target assignment was balanced between participants. Once the desired viscosity was reached, the tone began to play and participants were required to determine whether the tone was short or long compared to all tones they had heard so far (reference-free bisection) by moving the cursor to the corresponding target location on the right or left. If a response was made before the tone had elapsed, the trial was discarded and they were required to repeat the trial. No feedback was given. Viscosity and duration values were randomized across trials with equal representation in each block.

#### Experiment 2 (Temporal Reproduction)

In the second experiment, a separate group of participants performed a temporal reproduction task for tone intervals of 1000, 1500, 2000, 2500, 3000, 3500, and 4000 ms. Because the task had higher attentional demands and was more likely to cause fatigue, the 280 trials were segmented into 10 instead of five blocks. In this task, the manipulandum moved the participant’s arm (1000 ms) to the one of 16 encoding start locations in a grid-like configuration, and locked in place for 1000 ms until a green “go” cue appeared in the start location. Upon seeing the cue, participants were required to start moving. Moreover, the tone onset was contingent upon movement, and the trial was discarded and repeated if movement stopped before tone offset. After tone offset, a linearly ramping “brake” was applied to discourage further movement. Once movement stopped the manipulandum brought them to a central location for the reproduction phase. After seeing a green cue, they reproduced the heard interval by holding and releasing a button. No auditory or performance feedback was given. As in Experiment 1, duration and viscosity were randomized and equally represented in each block.

#### Analysis

In Experiment 1 and 2, movement length and force measures were taken for each trial. Movement length was defined as the summed distance traveled (point-by-point Euclidean distance between each millisecond time frame) during the stimulus tone. Force was similarly defined as the summed instantaneous force during the stimulus tone. In Experiment 1, reaction time (RT) was defined as the time elapsed between tone offset and reaching one of the two choice targets. Outlier trials were excluded for RT values greater than 3 standard deviations away from the mean of a participant’s log-transformed RT distribution (Ratcliff, 1993). For each participant we plotted duration by proportion of ‘long’ responses. From here, we used the psignifit software 4.0 package to estimate individual bisection points (BP) and coefficients of variation (CV) for all four viscosity values (Schütt et al., 2016); all curves were fit with a cumulative Gumbel distribution to account for the log-spaced nature of tested intervals (Wiener et al., 2018; Wiener et al., 2019).

In Experiment 2, we plotted true duration by estimated duration for each participant to find individual slope and intercept values. We also computed individual coefficient of variation (CV) values for duration and viscosity conditions via the ratio of estimation standard deviation to estimated mean (Wiener et al., 2019). We excluded trials if the reproduction time fell outside 3 standard deviations from the mean.

### Drift Diffusion Modeling

To better dissect the results of Experiment 1, we decomposed choice and RT data using a drift diffusion model (DDM; (Ratcliff, 1978; Wiecki et al., 2013; Tipples, 2015)). Due to the low number of trials available per condition, we opted to use hierarchical DDM (HDDM) as employed by the HDDM package for Python (http://ski.clps.brown.edu/hddm_docs/allinone.html). In this package, individual subjects are pooled into a single aggregate, which is used to derive fitted parameters by repetitive sampling from a hypothetical posterior distribution via Markov Chain Monte Carlo (MCMC) sampling. From here, the mean overall parameters are used to constrain estimates of individual-subject estimates. HDDM has been demonstrated as effective as recovering parameters from experiments with a low number of trials (Wiecki et al., 2013).

In our approach to modeling, we opted for a DDM in which four parameters were set to vary: the threshold difference for evidence accumulation (*a*), the drift rate towards each boundary (*v*), the starting point, or bias towards a particular boundary (*z*), and the non-decision time (*t*), accounting for remaining variance due to non-specific processes (e.g. perceptual, motor latencies). Our choice to include these four parameters was driven *a-priori* by earlier modeling efforts for studies of timing and time perception, in which all four parameters provide the best accounting for choice and RT (Tipples, 2015; Balcı and Simen, 2014; Wiener et al., 2018). Model construction was accomplished using the HDDMStimCoding class for HDDM, in which the starting point was split between both short and long response boundaries. Unlike the behavioral analysis, we included all trials here, and chose to model the probability of outliers using the *p outlier* option, in which outliers were assumed to come from a uniform distribution at the right tail of the full RT distribution (Ratcliff and Tuerlinckx, 2002). This was done to avoid differential weighting of RTs from individual subjects in the full distribution; the probability was set to 0.05. Model sampling was conducted using 10,000 MCMC samples, with a burn-in of 1000 samples and a thinning (retention) of every 5th sample. Model fits were assessed by visual inspection of the chains and the MC err statistic; all chains exhibited low autocorrelation levels and symmetrical traces. We additionally sampled 5 further chains of 5000 iterations (200 burn-in) and compared the Gelman-Rubin Statistic (Gelman and Rubin, 1992) revealing a value of 1.011 ± 0.082 (SD), indicating good chain stability.

In fitting our model, we chose to have all four free parameters vary by the viscosity presented on a given trial. We note that duration was not included as a variable, as the number of trials for each duration x viscosity combination was lower than recommended by HDDM (≤10). In a pilot analysis, we attempted to fit a larger model, but found large chain instability and error, indicating a lack of convergence.

#### Bayesian Observer Model

To model data from the reproduction task, we employed a Bayesian Observer Model, as developed by Jazayeri and colleagues (Jazayeri and Shadlen, 2010; Remington et al., 2018). In this model, sensory experiences of duration are treated as noisy estimates from a Gaussian distribution with scalar variability that grows linearly with the base interval, termed the measurement noise (*m*). Once drawn, these estimates are combined with the prior distribution of previously-experienced intervals; in this case, the prior was modeled as a uniform distribution with and upper and lower boundary corresponding to the presented intervals in the task. The mean of the resulting posterior distribution of an interval is thus drawn to the mean of the prior, thus accounting for the central tendency effect observed. Further, this effect also accounts for a trade-off in the precision of estimates; increased reliance on the prior, while increasing bias to the mean, also reduces variability, thus decreasing the CV (Cicchini et al., 2012). Following the posterior estimate, the produced movement is additionally corrupted by movement noise (*p*), again drawn from a Gaussian distribution. As an additional parameter, measurement bias is also included (*b*), also termed the estimation “offset” (Remington et al., 2018), in which the noisy estimate is shifted away from the true duration. Note here that *b* is specifically included as a shift in perception, rather than production bias.

Model parameters (*m, p, b*) were fit by minimizing the negative log-likelihood of individual subjects’ single trial responses, using modified code provided at (https://jazlab.org/resources/). Minimization was accomplished using the *fminsearch* function for Matlab, using numerical integration over the posterior distribution. Model fits were repeated using different initialization values and a fitting maximum of 3000 iterations; inspection of fitted parameters indicated good convergence of results.

## Acknowledgements

This work was supported by the National Science Foundation (1849067), awarded to MW and WMJ.

